# Detection and molecular characterization of a phytoplasma associated with Cucumber (*Cucumis sativus*) and its first report from India

**DOI:** 10.1101/2022.04.04.486938

**Authors:** Mantesh Muttappagol, H. D. Vinay Kumar, Shridhar Hiremath, M. Nandan, C. R. Jahir Basha, K. S. Shankarappa, V. Venkataravanappa, C. N. Lakshminarayana Reddy

## Abstract

Cucumber (*Cucumis sativus*) plants exhibiting typical phyllody symptoms were collected from farmers field of Chintamani, Chickballapur districts of Karnataka (India). Disease incidence of phyllody was 2-3%. The etiology of the cucumber phyllody phytoplasma (CuPP) was confirmed by amplifying 16S rRNA gene from symptomatic plants using PCR followed by nested PCR using universal primers pairs P1/P7 and R16F2n/R16R2. After general detection the non-ribosomal *Sec*Y and *rp* (ribosomal protein) genes was amplified using specific primers. The PCR amplified products 16SrRNA (1.2 kb), SecY (1.6 kb) rp gene (1.2 kb) was cloned and sequenced. Sequence and phylogenetic analyses of 16S rRNA, secY and *rp* genes revealed that the detected phytoplasma is a member of the 16SrI group (*Candidatus* Phytoplasma asteris). Further on the basis of computer-simulated RFLP (= *in silico* RFLP) analysis of amplified F2n/R2 region of 16S rRNA gene indicates that, the detected phytoplasma was belongs to the subgroup X (16SrII-X). This is the first report on phytoplasma associated with phyllody disease of cucumber in India.

## Introduction

Cucumber (*Cucumis sativus* L.) is an important vegetable crop belongs to family *Cucurbitaceae* grown widely in warm regions of the world. It is the world’s ancient vegetable crop originated in India (Yamaguchi 1983) and popularly known as *‘khira’* and gherkins. The crop is extensively cultivated for its edible tender fruit, preferred as salad ingredient, pickles and also as cooked vegetable (Navitha et al. 2019). In India cucumber is cultivated with in an area of 78,000 ha with the production of 1,142,000 MT (Anonymous, 2018 NHM data base). The cucumber is mainly gown during spring-summer and rainy season in commercial scale as well as kitchen gardens. The cultivation of cucumber is threaten by number insects and diseases from germination till harvest and causes heavy economic losses to the growers (Venkataravanappa et al. 2019).

Cucumber is reported to be attacked by plethora of viruses, bacteria and fungi in India, but not yet by phytoplasmas. All over the world, phytoplasmas cause a multitude of diseases in several hundred crop species including fruits, vegetables, cereals and trees (McCoy et al. 1989; Lee et al. 2000). Many cucurbits belongs to the *Cucurbitaceous* family have been infected with different phytoplasma has been well documented in different parts of the world that includes witches’ broom of *Cucurbita pepo* in India (Rao et al. 2017), cucumber phyllody in Iran (Hosseini et al. 2015) and cucumber yellows (Valiunas et al. 2017). Yellows of *C. maxima*, (McCoy et al., 1989), *C. moschata* (Montano et al. 2006), *Luffa cylindrica* L. (Lee et al. 1993; Gundersen et al. 1994), *Momordica charantia* L. (Alves et al. 2017), *Sechium edule* (Jacquin) Swartz (Villalobos et al. 2002), *Sicana odorifera* (Montano et al. 2007), *Trichosanthes cucumerina* (Weng et al. 2021) and *C. moschata* L., (Xu et al. 2021).The phytoplasma is one of the emerging pathogens in many vegetable crops and causing huge loss to various crops. Based on the 16Sr RNA gene analysis, 16 different ribosomal groups and subgroups phytoplasmas infecting different vegetable crops have been well documented across the world (Kumari et al. 2019). Considering nutritional benefits of cucumbers in human diet, the crop is protected against biotic stress. With this backdrop, the present was attempted to identify and characterize strain of the phytoplasma associated phyllody disease of cucumber based on PCR assays and 16S rRNA, *Sec*Y and rp gene sequence comparison analysis.

## Materials and Methods

### Collection of disease samples

During April 2021, typical phytoplasma like symptoms were observed in cucumber in different farmers fields in Chintamani (13.40°N 78.06°E.), Chickballapur (Dist.) Karnataka (India). The infected cucumber plants displayed typical symptoms of phytoplasma infection such as witches’ broom, flower viresence, small leaves, and dense clusters of highly proliferating apical shoot. Disease incidence was calculated as the percentage of symptomatic plants over total number of plants observed in an area of 20 × 20 meter plot. Due to creeping growth habit of cucumber, care was taken not to sample the same plant more than once, total five symptomatic and one symptomatic leaf samples were collected from same location. A part of the collected leaf samples were used for DNA isolation and the remaining sample was stored at −80°C for further use.

### DNA extraction and PCR amplification of 16SrRNA, *SecY* gene, and ribosomal protein (rp) gene

Total genomic DNA of symptomatic and asymptomatic cucumber plants was extracted using cetyl-trimethyl ammonium bromide (CTAB) method. (Dolye and Doyle 1990). The DNA extracted from the phytoplasma infected bitter gourd (16SrI, KX179474) and bottle gourd (16SrI, MT386510) was included as positive control in PCR assays (Venkataravanappa et al. 2017; Ashwathappa et al. 2020). Using universal 16Sr RNA gene specific primers pairs (P1/P7, (R16mF2/R16mR2), the phytoplasma infection in cucumber was confirm in both direct and nested PCR assays (Deng and Hiruki 1991; Schneider 1995); (Gundersen and Lee 1996).The 16S rRNA gene sequence is highly conserved and insufficient for finer level differentiation of phytoplasma strains, therefore housekeeping genes such as translocase subunit *sec*Y and ribosomal protein (*rp*) was amplified by PCR using primer pairs (SecYF1/SecYR1) (Lee et al. 2010) and (*rp*(II)F1A/*rp*(II)R1A) (Martini et al., 2013). PCR reactions were carried out in a thermal cycler (Master cycler, Profelex, Hamsburg, Germany) and the cycling protocol used for PCR assay was followed as earlier described (Venkataravanappa et al. 2017).Then five microliters of PCR products were analysed in 0.8% (W/V) ethidium bromide stained agarose gel, and visualized with a UV transilluminator. The amplified PCR products of 16SrRNA, *Sec*Y and *rp* genes were purified from agarose gel using Qiaquick gel extraction kit (Qiagen, Hilder, USA) and cloned into pTZ57R/T cloning vector according to manufacturer’s directions (MBI Fermentas, USA). The transformation was performed using *Escherichia coli* (DH5α) cells. The recombinant plasmids were purified and confirmed with restriction digestion (*Pst*1 and *Sac*1 enzymes). The confirmed clones were sequenced in both directions at Eruofins Genomics Pvt. Ltd., Bengaluru, India.

### Sequence analysis

The full-length sequences of 16S rRNA, *secY* gene and *rp* genes obtained in the present study were subjected to BLAST search for finding the similar sequences in the database. The sequences showing maximum homology with 16SrRNA (Table 1), *secY* gene (Table 2) and *rp* gene (Table 3) of cucumber phyllody phytoplasma were retrieved from NCBI database and aligned using BioEdit (Hall 1999) and ClustalW (Thompson et al. 1997) program. MEGA X software was used for constructed phylogeny using Neighbour-Joining method with 1000 bootstrapped replications (Kumar et al. 2016). Sequence of *Acholeplasma laidlawii* (Accession number NR076550) for 16S rRNA, *Leptospira interrogans* (Accession number MH376290) for *secY* gene sequences and *Acholeplasma laidlawii* (Accession number M81465) were used as out groups in the phylogenetic tree construction.

**Table:1.** 16Sr RNA gene sequences of phytoplasmas employed in analysis.

| Phytoplasma disease / country | 16S rDNA group/subgroup | GenBank accession number | Identity with Cucumber phytoplasma |
| --- | --- | --- | --- |
| Cucumber phyllody phytoplasma/ Iran | 16SrII-D | KR822804 | 67.30 |
| Cucumis sativus' phyllody/Iran | 16SrVI | KR633068 | 61.90 |
| Cucumber phyllody phytoplasma/Iran | 16SrII | JN574839 | 72.20 |
| '*Candidatus* Phytoplasma trifolii'/Iran | 16SrVI | KT807467 | 62.20 |
| Cucumber viresence /Taiwan | 16SrI | GU361755 | 99.10 |
| Aster yellows/USA | 16SrI-B | M30790 | 68.60 |
| Potato purple top/Russia | 16SrI-B | EU333396 | 68.60 |
| Potato purple top/Russia | 16SrI-P | EU333397 | 68.60 |
| Apricot aster yellows/Spain | 16SrI-F | AY265211 | 68.30 |
| Aconitum proliferation/ Lithuania | 16SrI-A | AF510323 | 68.40 |
| Aster yellows-AV2192/Germany | 16SrI-L | AY180957 | 68.60 |
| Aster yellows-AVUT/Germany | 16SrI-M | AY265209 | 68.70 |
| Rhododendron x hybridum/ Czech | 16SrI-C | KC009838 | 68.40 |
| Republic |  |  |  |
| Aster yellows-BB/USA | 16SrI-A | AY180955 | 68.40 |
| Blueberry stunt-BBS3/USA | 16SrI-E | AY265213 | 68.40 |
| Aster yellows Btsv2CarD3/USA | 16SrI-B | AY180945 | 68.60 |
| Cherry little leaf/ Lithuania | 16SrI-Q | AY034089 | 68.50 |
| Japanese maple witches' broom/ | 16SrI-B | JQ015183 | 68.70 |
| Clover phyllody/Germany | 16SrI-C | AY265217 | 68.40 |
| Valeriana yellows/ Lithuania | 16SrI-M | AY102274 | 68.10 |
| Aster yellows / USA | 16SrI-O | AF268405 | 68.70 |
| *Cucumis sativus*' stunt virescence/<br>Lithuania | 16SrI-A | KX298483 | 68.40 |
| *Cucumis. sativus*' phytoplasma /Iran | 16SrVI-A | MH004460 | 62.50 |
| *Cucumis sativus*' phytoplasma /Iran | 16SrVI-A | MH004459 | 39.40 |
| Western X-disease/USA | 16SrIII-A | L04682 | 61.90 |
| Coconut lethal yellowing/USA | 16SrIV-A | U18747 | 62.20 |
| Elm yellows/USA | 16SrV-A | AY197655 | 62.20 |
| Jujube witches' broom/Korea | 16SrV-B | AB052876 | 61.30 |
| "Flavescence dorée/France | 16SrV-C | AF176319 | 62.20 |
| Clover proliferation/Canada | 16SrVI-A | AY390261 | 62.20 |
| Ash yellows/USA | 16SrVII-A | AF092209 | 62.40 |
| Apple proliferation/ Germany | 16SrX-A | AJ542541 | 63.40 |
| Spartium witches' broom/Italy | 16SrX-B | X92869 | 63.50 |
| Pear decline/ Germany | 16SrX-C | AJ542543 | 63.60 |
| European stone fruit yellows/<br>Germany | 16SrX-F | AJ542544 | 63.50 |
| Rice yellow dwarf/Germany | 16SrXI-A | AB052873 | 60.80 |
| *Cucumis sativus* phytoplasma /Iran | 16SrVI-A | MH004458 | 39.40 |
| Squash phyllody phytoplasma/Iran | 16SrII-D | KR822805 | 67.40 |
| Strawberry yellows / Lithuania | 16SrXII-E | DQ086423 | 66.80 |
| Mexican periwinkle<br>virescence/Mexico | 16SrXIII-A | AF248960 | 66.40 |
| Bermudagrass white leaf/Italy | 16SrXIV-A | AJ550984 | 62.00 |
| Chestnut witches' broom/South<br>Korea | 16SrXIX-A | AB054986 | 61.40 |
| Hibiscus witches' broom/Brazil | 16SrXV-A | AF147708 | 61.50 |
| Sugarcane yellow leaf/Cuba | 16SrXVI-A | AY725228 | 64.80 |
| American potato purple top wilt/USA | 16SrXVIII-A | DQ174122 | 66.10 |
| Buckthorn witches' broom/USA | 16SrXX-A | X76431 | 63.10 |
| Pine shoot proliferation/Europe | 16SrXXI-A | AJ632155 | 62.00 |
| Awka wilt/Nigeria | 16SrXXII-A | Y14175 | 61.80 |
| Buckland Valley grapevine yellows/<br>Australia | 16SrXXIII-A | AY083605 | 66.70 |
| Sorghum bunchy shoot/USA | 16SrXXIV-A | AF509322 | 59.20 |
| *Cassia italica* witches' broom/Oman | 16SrXXIX-A | EF666051 | 61.40 |
| Weeping tea witches'<br>broom/Australia | 16SrXXV-A | AF521672 | 61.20 |
| Sugarcane yellows D3T1/ Mauritius | 16SrXXVI-A | AJ539179 | 64.40 |
| Salt cedar witches' broom/China | 16SrXXX-A | FJ432664 | 63.80 |

**Table:2.** *SecY* gene sequences of phytoplasmas employed in the analyses.

| Phytoplasma Species /Country | Sub-group | GenBank<br>accession No. | Identity with<br>Cucumber phytoplasma |
| --- | --- | --- | --- |
| Tomato big bud phytoplasma/USA | 16SrI-A | AY803178 | 83.40 |
| Chrysanthemum yellows phytoplasma/ Germany | 16SrI-A | AY803170 | 83.20 |
| Hydrangea phyllody phytoplasma/Belgium | 16SrI-A | AY803181 | 83.20 |
| Chrysanthemum yellows phytoplasma/Italy | 16SrI-B | DQ787851 | 87.80 |
| *Trema levigatum*' witches'-broom/China | 16SI-B | MW032211 | 98.75 |
| Primrose virescence phytoplasma/Germany | 16SrI-B | AY803176 | 86.80 |
| Clover phyllody phytoplasma/Germany | 16SrI-C | AY803183 | 83.80 |
| Paulownia witches-broom phytoplasma/Taiwan | 16SrI-D | AY803184 | 86.50 |
| Blueberry stunt phytoplasma/ Michigan, USA | 16SrI-E | AY803169 | 84.20 |
| Apricot chlorotic leaf roll phytoplasma/Spain | 16SrI-F | AY803166 | 83.50 |
| Strawberry multiplier phytoplasma/Florida, USA | 16SrI-K | AY803180 | 83.50 |
| Aster yellows phytoplasma/USA | 16SrI-M | AY803168 | 86.60 |
| Ipomoea witches-broom phytoplasma/Taiwan | 16SrI-N | AY803182 | 98.58 |
| Peanut witches-broom phytoplasma/Taiwan | 16SrII-A | GU004331 | 26.80 |
| Sesame phyllody phytoplasma/ Thailand | 16SrII | GU004322 | 28.40 |
| Sesame phyllody phytoplasma/ Thailand | 16SrII | GU004362 | 37.50 |
| Soybean phyllody phytoplasma/ Thailand | 16SrII-C | GU004324 | 28.50 |
| Picris echioides phytoplasma/Italy | 16SrII-E | GU004348 | 37.80 |
| Australian tomato big bud phytoplasma/ Australia | 16SrII-D | GU004347 | 37.70 |
| Sesame phyllody phytoplasma/ Myanmar | 16SrII | AB703253 | 37.80 |
| Milkweed yellows phytoplasma/ New York, USA | 16SrIII-F | GU004340 | 30.00 |
| Potato purple top phytoplasma/ Montana, USA | 16SrIII-M | GU004333 | 29.90 |
| Clover yellow edge phytoplasma/ Lithuania | 16SrIII-B | GU004332 | 30.30 |
| Spirea stunt phytoplasma/ New York, USA | 16SrIII-E | GU004326 | 28.20 |
| Poinsettia branch-inducing phytoplasma/USA | 16SrIII-H | GU004328 | 30.00 |
| Peach X-disease phytoplasma/Canada | 16SrIII-A | GU004327 | 28.10 |
| Walnut witches-broom phytoplasma/ Georgia, USA | 16SrIII-G | GU004325 | 30.00 |
| Elm yellows phytoplasma/ New York, USA | 16SrV-A | AY197690 | 41.00 |
| Cherry lethal yellows phytoplasma/China | 16SrV-B | AY197693 | 41.30 |
| Alder yellows phytoplasma/Germany | 16SrV-C | AY197692 | 41.20 |
| Rubus stunt phytoplasma/Italy | 16SrV-E | AY197696 | 41.50 |
| Flavescence doree phytoplasma/USA | 16SrV-D | AY197685 | 41.40 |
| Dry bean phyllody phytoplasma/ Washington, USA | 16SrVI-A | GU004353 | 40.30 |
| Ash yellows phytoplasma/ NY California, USA | 16SrVI-A | GU004329 | 28.60 |
| Brinjal little leaf phytoplasma/India | 16SrVI-D | GU004356 | 30.30 |
| Potato witches'-broom phytoplasma/Canada | 16SrVI-A | GU004316 | 29.00 |
| Clover phyllody phytoplasma/Canada | 16SrVI-A | GU004315 | 29.10 |
| Potato purple top phytoplasma / Alaska, USA | 16SrVI-A | GU004343 | 29.20 |
| Lucerne virescence phytoplasma/Frans | 16SrVI-A | GU004318 | 30.00 |
| Vinca virescence phytoplasma/ California, USA | 16SrVI-A | GU004317 | 28.70 |
| Potato purple top phytoplasma/ Alaska, USA | 16SrVI-A | GU004344 | 37.30 |
| Potato purple top phytoplasma/ Alaska, USA | 16SrVI-A | GU004342 | 38.20 |
| Potato purple top phytoplasma/ Alaska, USA | 16SrVI-A | GU004351 | 38.70 |
| Dry bean phyllody phytoplasma/USA | 16SrVI-A | GU004352 | 40.00 |
| Coconut lethal yellows phytoplasma/USA | 16SrIV-A | GU004320 | 29.60 |
| *Candidatus* Phytoplasma fraxini/USA | 16SrVII-A | GU004329 | 28.60 |
| Apple proliferation phytoplasma/Italy | 16SrX-A | GU004335 | 29.60 |
| Pear decline phytoplasma/Italy | 16SrX-C | GU004363 | 36.60 |
| Mexican periwinkle virescence /Meixco | 16SrXIII-A | GU004336 | 29.10 |
| American potato purple top wilt phytoplasma/USA | 16SrXVIII-B | GU004338 | 29.50 |
| Stolbur-It phytoplasma/Italy | 16SrXII-A | GU004355 | 47.50 |
| '*Candidatus* Phytoplasma americanum'/USA | 16SrXVIII-B | MN227134 | 38.00 |

**Table.3.** Rep gene sequences of phytoplasmas employed in analysis.

| Phytoplasma Species/ Country | Sub-group | GenBank accession No. | Identity with Cucumber phytoplasma |
| --- | --- | --- | --- |
| *Gynostemma pentaphyllum*' witches'-broom/China | 16SrI-B | MN543070 | 99.00 |
| Aster yellows phytoplasma/Hungary | 16SrI-B | MN526023 | 99.10 |
| Rapeseed phyllody/Taiwan | 16SrI | CP055264 | 99.30 |
| Plum witches'-broom phytoplasma/Poland | 16SrI-B | MH061366 | 96.10 |
| Oilseed rape phytoplasma/China | 16SrI-A | KX551965 | 99.10 |
| *Melochia corchorifolia*' phyllody/China | 16SrI-A | KX158198 | 99.10 |
| *Tilia platyphyllos*' phytoplasma/ Lithuania | 16SrI-B | JX184926 | 97.90 |
| *Salix tetradenia*' witches'-broom/China | 16SrI-B | KC117314 | 99.10 |
| Periwinkle phyllody/Korea | 16SrI-B | AB741642 | 98.20 |
| Sesame phyllody/Korea | 16SrI-L | AB741638 | 97.70 |
| Larch phytoplasma strain / Lithuania | 16SrI-B | JF767009 | 98.60 |
| Periwinkle phyllody/China | 16SrI-B | HQ285145 | 99.20 |
| Iranian tomato phytoplasma /Iran | 16SrI-B | HQ286478 | 99.10 |
| Onion proliferation phytoplasma/ Lithuania | 16SrI-A | GU228514 | 96.90 |
| Aster yellows/USA | - | AY183710 | 96.90 |
| Paulownia witches'-broom/Germany | - | AY264857 | 96.80 |
| Aster yellows phytoplasma/Germany | - | AY183708 | 96.10 |
| Green gram phyllody/ Myanmar | 16SrII | AB703244 | 96.70 |
| Black gram witches' broom/Myanmar | 16SrII | AB703243 | 57.10 |
| Parthenium weed witches' broom/ Yunnan | 16SrII | KU958723 | 57.10 |
| Sword bean witches' broom/ Yunnan | 16SrII | KC953009 | 57.70 |
| Peanut witches' broom/ Yunnan | 16SrII | JX871469 | 57.60 |
| Cucumber phyllody/Iran | 16SrII | KY365523 | 57.50 |
| Sesame phyllody/ Myanmar | 16SrII | AB703247 | 56.80 |
| Peach X-disease/USA | 16SrIII-A | EF186813 | 55.60 |
| Goldenrod yellows/USA | 16SrIII-D | EF186810 | 57.80 |
| Vaccinium witches' broom/ Germany | 16SrIII-F | EF186809 | 57.70 |
| Poinsettia branch inducing/ USA | 16SrIII-H | EF186811 | 58.30 |
| Gliricidia little leaf/ Honduras | 16SrIX | EF186800 | 57.80 |
| Knautia arvensis phyllody/ Italy | 16SrIX | EF186801 | 60.40 |
| Pigeon pea witches' broom/USA | 16SrIX-A | EF183495 | 60.20 |
| Almond witches' broom/ Lebanon | 16SrIX-B | EF186803 | 59.70 |
| Picris echioides phyllody/Italy | 16SrIX-C | EF186802 | 59.90 |
| Elm yellows/USA | 16SrV-A | AY197675 | 59.40 |
| Alder yellows/Italy | 16SrV-C | AY197666 | 59.00 |
| Spartium witches' broom/Italy | 16SrV-C | AY197672 | 58.60 |
| Hemp dogbane yellows/USA | 16SrV-C | AY197674 | 58.20 |
| "Flavescence dorée"/Italy | 16SrV-D | AY197664 | 58.80 |
| Catharanthus phyllody/Sudan | 16SrVI | EF183494 | 58.60 |
| Ash yellows/USA | 16SrVII-A | EF183492 | 58.50 |
| Apple proliferation/Italy | 16SrX-A | EF193366 | 58.60 |
| Pear decline/Germany | 16SrX-C | EF193370 | 67.60 |
| Mexican periwinkle virescence/Mexico | 16SrXIII-A | EF193365 | 67.70 |
| American potato purple top wilt/USA | 16SrXVIII-A | EF193362 | 77.90 |
| Tomato "stolbur"/Italy | 16SrXII-A | EF193364 | 78.00 |
| Palm lethal yellowing/Florida | 16SrIV-A | DQ297677 | 80.10 |
| Coconut lethal yellowing/ Honduras | 16SrIV-A | DQ318238 | 47.40 |
| Texas Phoenix palm phytoplasma/USA | 16SrIV-A | EF050087 | 47.30 |
| Lethal yellowing phytoplasma/Jamaica | 16SrIV-A | EF186805 | 47.50 |
| Lethal yellowing phytoplasma/USA | 16SrIV-A | EF186804 | 60.60 |
| Coconut lethal yellowing/USA | 16SrIV-A | EF193382 | 60.50 |
| Mexican periwinkle virescence/ Mexico | 16SrXIII | EF193365 | 60.30 |
| '*Candidatus* Phytoplasma meliae'/Argentina | 16SrXIII | KU850947 | 77.90 |
| '*Candidatus* Phytoplasma meliae'/South Korea | 16SrXIII | KU850945 | 78.20 |
| '*Candidatus* Phytoplasma fragariae'/USA | 16SrXIII | KJ144893 | 78.20 |

### Virtual RFLP analysis

Virtual RFLP (Restriction fragment length polymorphism) analysis was carried out for F2n/R2 region of the 16Sr RNA gene derived from cucumber phyllody phytoplasma using *iPhy*Classifier online tool (https://plantpathology.ba.ars.usda.gov/cgi-bin/resource/iphyclassifier.cgi) (Zhao et al. 2009) to identify the phytoplasma to group and subgroup level. The virtual RFLP patterns were compared, and the similarity coefficient was calculated (Wei et al. 2008). Restriction pattern obtained for the phytoplasmas associated with cucumber phyllody was compared with already published RFLP patterns of phytoplasma representatives of available ribosomal groups and 16SrI subgroups (Lee et al. 1998).

## Results

The predominant symptoms observed on phytoplasma infected cucumber plants are little leaf, excessive proliferation along the stem, flower virescence and phyllody, internode shortening phyllody and witches’ broom (Fig1a-d). These symptoms are similar to those described for cucumber phyllody disease reported to occur in other countries (Salehi et al. 2015). The cucumber phyllody disease incidence was ranged from 2-5%.

**Fig: 1.**
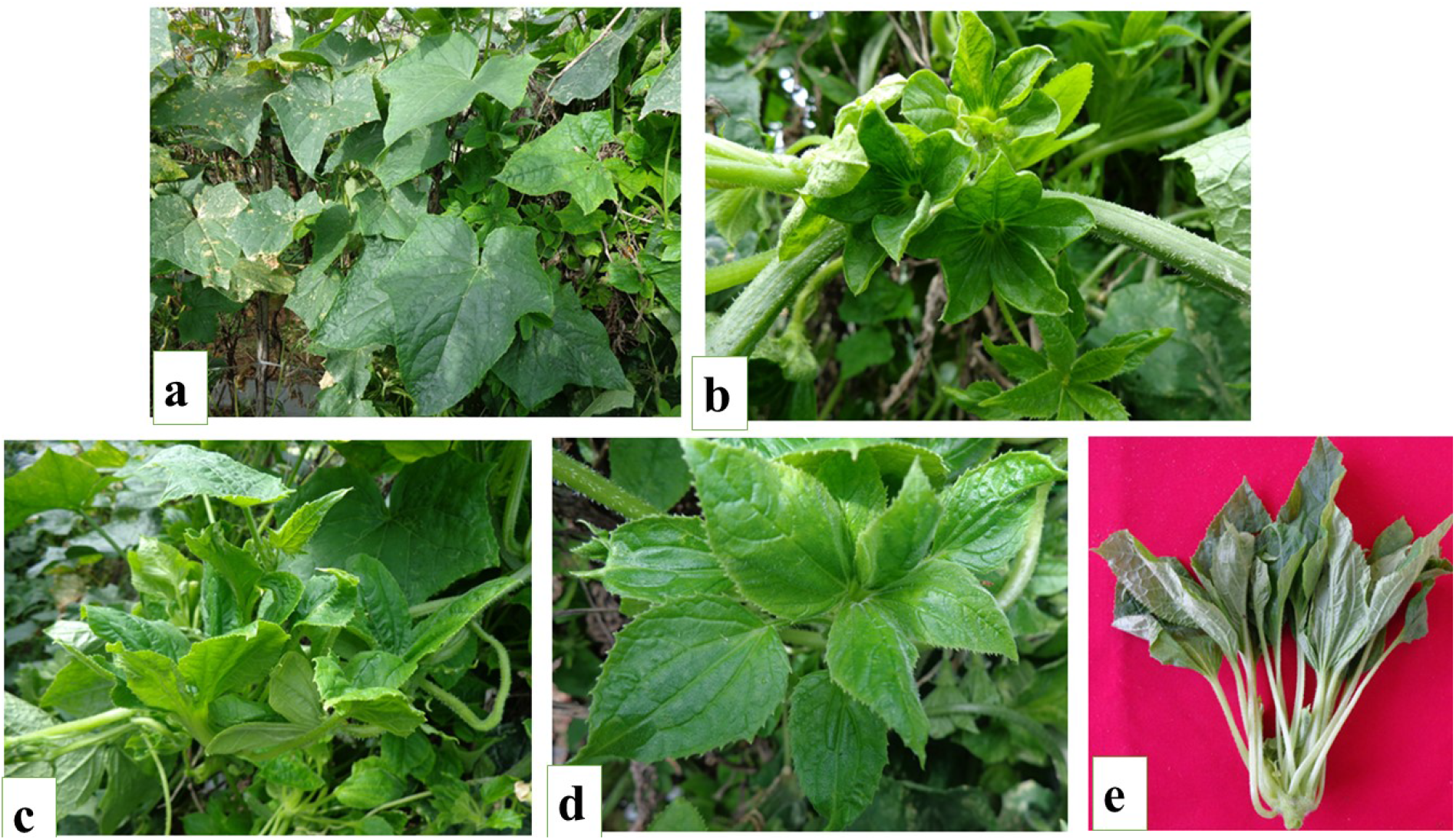
Symptoms on cucumber (*Cucumis sativus*) plant var. Rashmi under field condition **a**; flower virescence with phyllody **b**; excessive shoot proliferation **c**; Little leaf **d**; cucumber plant with floral virescence and small leaves, internode shortening **e**.

The resulting PCR amplicon of 1.2 kb corresponding to the phytoplasma 16Sr RNA gene, *secY* gene (1.6 kb) and *rp* gene (1.2 kb) were amplified in all four cucumber samples displaying phyllody disease symptoms and positive control samples. However, no such amplification was observed in asymptomatic plants with each of primer pair employed. Amplified PCR fragments of each gene was cloned separately and sequenced in both the direction. The consensus sequences of 16S rRNA (accession number MZ401356), *secY* gene (accession number MZ463022) and *rp* gene (accession number MZ463023) were deposited in GenBank.

BLASTn searches for homologous gene sequences of 16S rRNA gene sequence of cucumber phyllody isolate obtained in the study was more homologous to those of phytoplasmas classified in the, 16SrI group (‘*Candidatus* Phytoplasma asteris’) having the maximum nucleotide(nt) sequence identity of 99.13% with cucumber viresence from Taiwan (GU361755). The result was well supported in phylogenetic analysis of 16S rRNA gene with selected reference strains indicated that the cucumber phyllody phytoplasma is clustered together with several strains of phytoplasma belongs to the 16SrI subgroup (Fig. 2a).

**Figure: 2.**
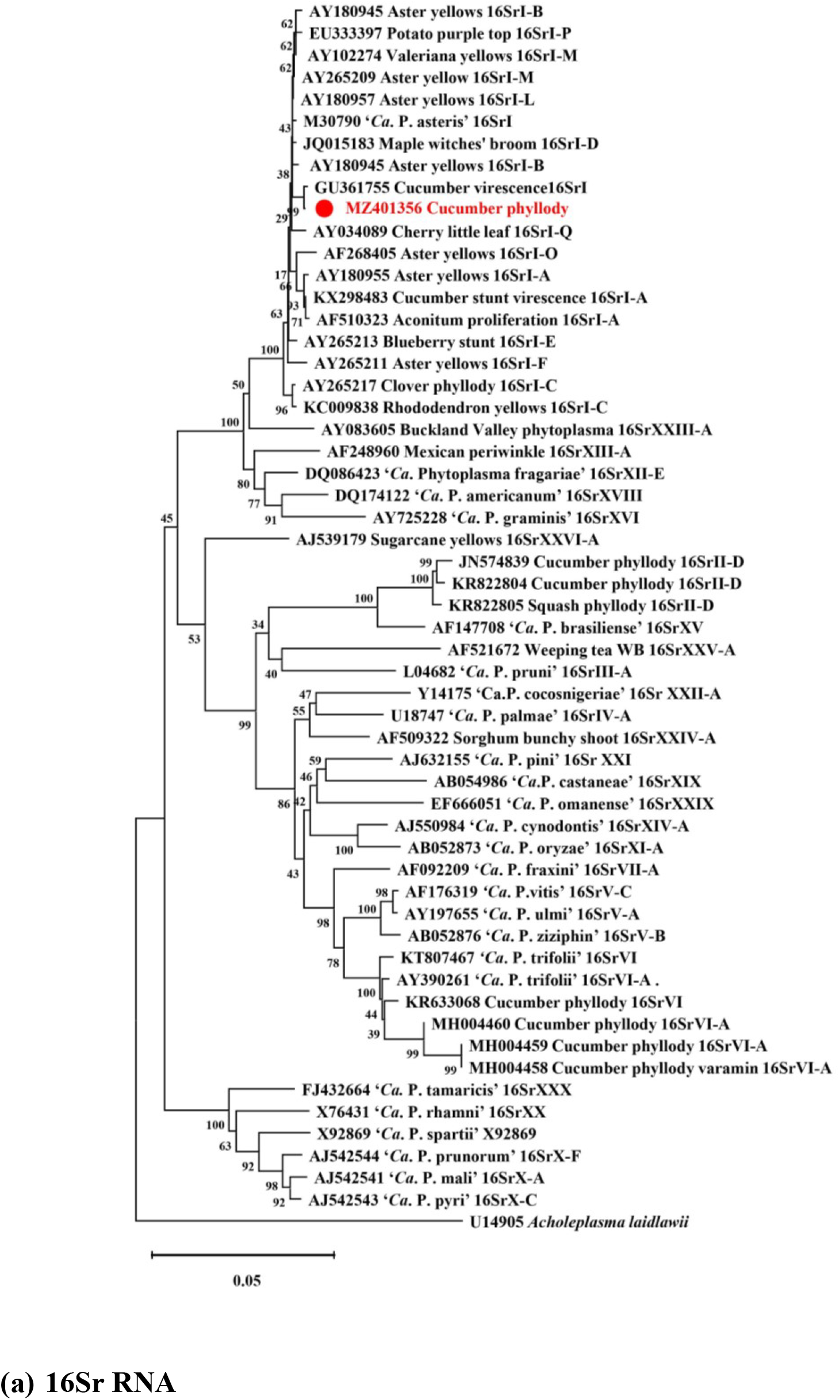
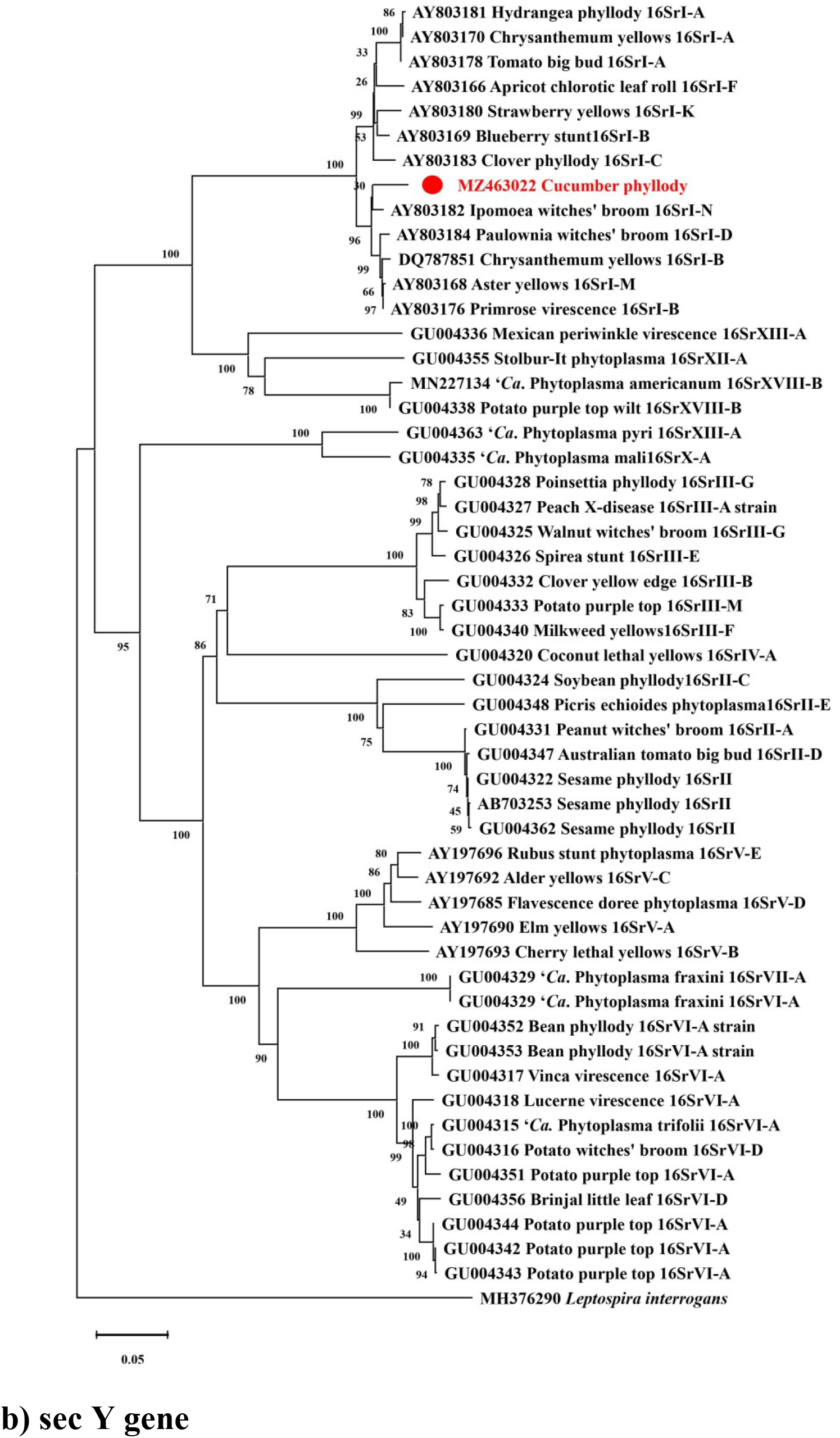
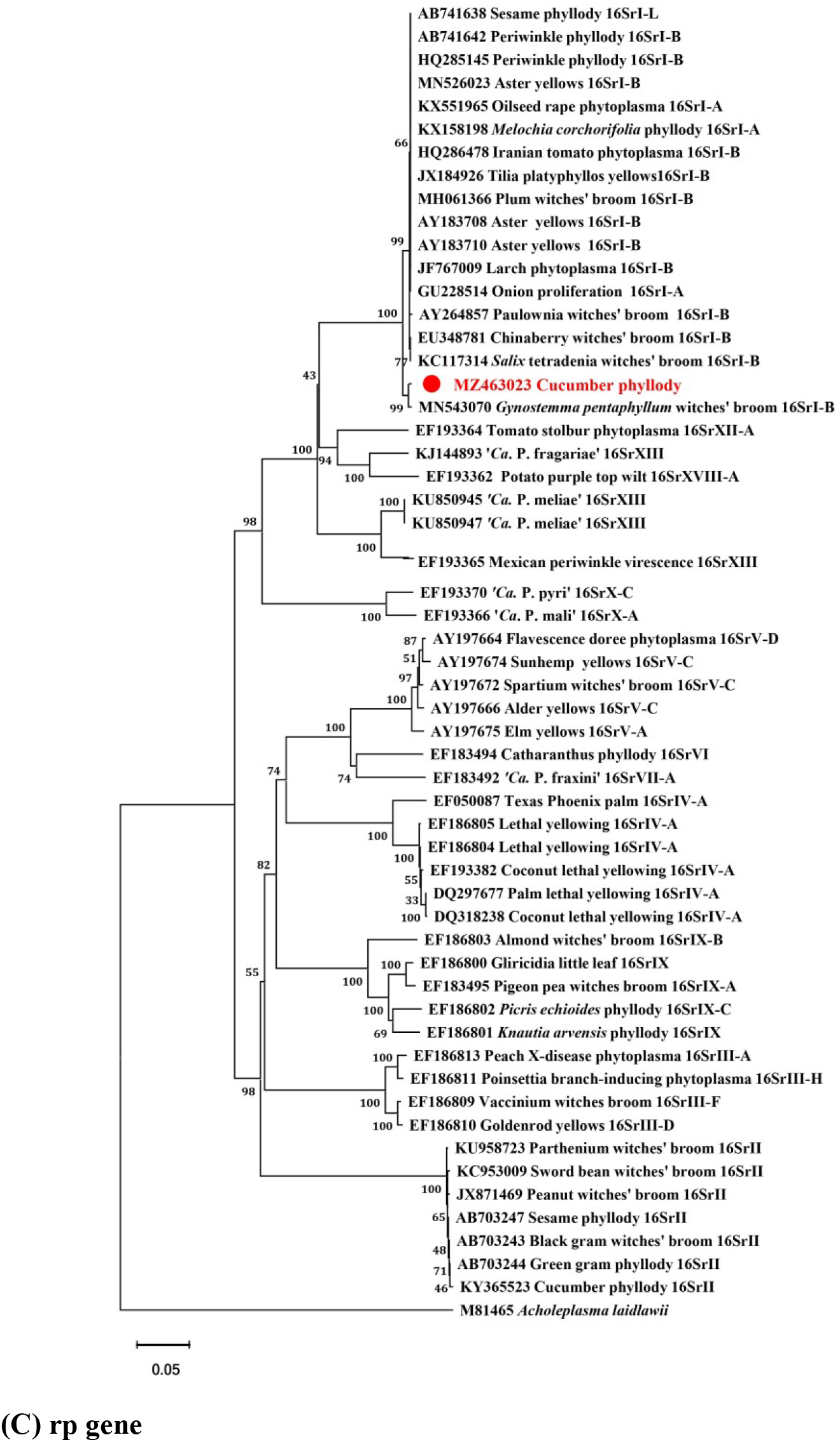
Phylogenetic tree constructed with MEGA X Software by utilizing 16S rRNA **(a)** partial sec *Y* **(b)** and partial rp **(c)** gene sequences of cucumber phyllody phytoplasma (in red) with phytoplasma strains listed in Tables 1, 2, and 3. The tree was rooted with *Acholeplasma laidlawii* (GenBank accession number U14905), *Leptospira interrogans* (GenBank accession number MH376290) and *Acholeplasma laidlawii* (GenBank accession number M81465) respectively. A bootstrap analysis with 1,000 replicates was performed and the bootstrap percent values above 50 are showed along the branches.

The computer-simulated RFLP (*in silico* RFLP) analysis of F2n/R2 region derived from 16S rRNA gene sequence of cucumber phyllody phytoplasma was identical (similarity coefficient 0.98) to the reference pattern of 16Sr group I, subgroup X (GenBank accession number JF781308). Therefore, cucumber phyllody phytoplasma having similarity coefficient more than the threshold level (Wei et al. 2008) and it may be considered as a variant of 16SrI-X subgroup under “*Candidatus* Phytoplasma asteris” 16Sr I group.

The *secY* sequence of cucumber phyllody phytoplasma showed the maximum nt similarity of 98.75% to that of *Trema levigatum’* witches’-broom from China (GenBank accession number MW032211). Similarly, *rp* gene of cucumber phyllody phytoplasma shared maximum nt similarity of 99.34% to that rapeseed phyllody phytoplasma from Poland (accession number CP055264). This result was also well supported by the phylogenetic analysis, in which *secY* (MZ463022) and *rp* (MZ463023) gene sequences of the cucumber phyllody phytoplasma is closely cluster with several phytoplasmas belongs to *Candidatus* Phytoplasma asteris” 16Sr-I group (Fig. 2b and 2c).

Attempt to detect phytoplasma in the cucumber samples by LAMP assay using different LAMP primers specific to 16S RNA gene showed that CuP was successfully detected in the infected cucumber samples (Fig. 7a). No amplification was observed from the DNA of negative sample and water control. The LAMP amplicons (a ladderlike pattern) were observed on 1.5% agarose gel from positive samples, but not from the negative sample and water control. The positive reactions can be visualized directly by the naked eye with samples showing sky blue color pattern, whereas a violet color change is observed in negative and water control (Fig. 7b).

Phytoplasma diseases cause significant economic losses in numerous *Cucurbitaceae*-related vegetable crops, including *C. maxima* (Giant pumpkin), *C. mixta* (winter Squash), *C. pepo (summer squash)* (Rao et al. 2017), *Lagenaria leucantha* (Bottle gourd) (Ashwathappa et al. 2020; Tripathi et al. 2017), *C. moschata* (Butternut Pumpkin) (Montano et al. 2006; Xu et al. 2021), *Momordica charantia* (Bitter gourd) (Montano et al. 2007; Venkataravanappa et al. 2017). However, there is no information is available for phytoplasma infection in cucumber in India. Literature survey showed that cucumber acts as natural host for different phytoplasma groups, that includes 16SrII-M from Iran (Dehghan et al. 2015; Ghayeb Zamharir and Azimi 2018), 16SrVI-A (Salehi et al. 2015) from Iran. In the present study the cucumber phyllody phytoplasma resulted from association of 16SrI group phytoplasma was confirmed based 16Sr RNA gene characterization including *secY* and *rp* genes. This study represents the first recorded of phytoplasma association with cucumber in Karnataka (India) is related to ‘*Ca*. P. asteris’-related strain belonging to 16Sr I-X subgroup. This disease become major threat for cultivation of cucumber as it causes 100% yield losses due to shoot and floral malformation and sterility. Moreover, the crop is a regular and widely cultivated throughout India in large farming community, therefore, further studies on epidemiology along with possible natural spread sources and management is also a foremost need.

## Contributions

MM, HDV and MN performed the biological experiment, SH, VV and KSS involved in the bioinformatics analysis CRJB and CNL involved in experimental design, CNL conceptualized work and provided the overall direction, all authors are involved in literature mining and manuscript preparation. All authors read and approved the final manuscript

## Ethics declarations

This article does not contain any studies requiring ethical approval.

## Conflict of interests

The authors declare that they have no competing interests.

## Human and animal rights

This article does not contain any studies with human or animal subjects performed by any of the authors.

## Informed consent

Informed consent was obtained from all individual participants included in the study.

## Notes

### Competing Interest Statement

The authors have declared no competing interest.

### Summary of Updates

Authors list has been updated

